# Robust cone-mediated signaling persists late into rod photoreceptor degeneration

**DOI:** 10.1101/2022.04.27.489758

**Authors:** Miranda L. Scalabrino, Mishek Thapa, Lindsey A. Chew, Esther Zhang, Jason Xu, A.P. Sampath, Jeannie Chen, Greg D. Field

**Affiliations:** Department of Neurobiology, Duke University School of Medicine, Durham, NC; Department of Statistical Science, Duke University, Durham NC; Jules Stein Eye Institute, University of California Los Angeles, Los Angeles, CA.; Zilkha Neurogenetics Institute, Keck School of Medicine, University of Southern California, Los Angeles, CA.

## Abstract

Rod photoreceptor degeneration causes deterioration in the morphology and physiology of cone photoreceptors along with changes in retinal circuits. These changes could diminish visual signaling at cone-mediated light levels, thereby limiting the efficacy of treatments such as gene therapy for rescuing normal, cone-mediated vision. However, the impact of progressive rod death on cone-mediated signaling remains unclear. A mouse model of rod degeneration was used to investigate the fidelity of retinal ganglion cell (RGC) signaling throughout disease progression. Despite clear deterioration of cone morphology with rod death, cone-mediated signaling among RGCs remained surprisingly robust: spatiotemporal receptive fields changed little and the mutual information between stimuli and spiking responses was relatively constant. This relative stability held until nearly all rods had died and cones had completely lost well-formed outer segments. Interestingly, RGC information rates were higher and more stable for natural movies than checkerboard noise as degeneration progressed. The main change in RGC responses with photoreceptor degeneration was a decrease in response gain. These results suggest that gene therapies for rod degenerative diseases are likely to successfully prolong cone-mediated vision even if there are changes to cone morphology and density.

## Introduction

Rod photoreceptor degeneration frequently leads to cone photoreceptor degeneration and death. Well before all rods have died, cones exhibit clear changes in morphology and physiology (Hartong, Berson, & Dryja, 2006; Sahel, Bonnel, Mrejen, & Paques, 2010). This is likely due to the loss of trophic factors and structural support provided by rods to nearby cones (Campochiaro & Mir, 2018). The extent to which these changes in cone structure and function compromise the ability of the retina to reliably signal visual scenes at high light levels is not clear. One possibility is that rod degeneration has an immediate impact on the ability of the retina to reliably signal visual scenes at cone-mediated light levels. Alternatively, recent work has highlighted homeostatic mechanisms in retinal circuits that help compensate for photoreceptor loss (Care et al., 2020; Lee, Care, Santina, & Dunn, 2021; Shen, Wang, Soto, & Kerschensteiner, 2020). Such mechanisms could facilitate reliable signaling at the level of retinal output, despite deterioration in photoreceptor function. Resolving these possibilities will inform treatment timepoints for gene therapies aimed at halting rod loss to preserve cone-mediated vision.

To examine the impact of progressive rod loss on cone-mediated visual signaling, we used a mouse line that models a human form of retinitis pigmentosa (RP), a blinding disorder characterized by initial degeneration of rod photoreceptors that ultimately leads to cone degeneration. This mouse line, *Cngb1^neo/neo^*, contains an insertion that interrupts translation of Cngb1, the beta subunit of the cyclic nucleotide gated cation channel in rods (Chen et al., 2010). Without this subunit, normal channels fail to form, causing rods to be tonically hyperpolarized and ultimately resulting in rod death (Hüttl et al., 2005; Zhang et al., 2009). This degeneration is relatively slow, with approximately 30% rod loss at 1 month (M) postnatal, complete loss of rods by 7M postnatal, and complete cone loss by 8-9M postnatal. Slow degeneration in this model provides a relatively large temporal window in which to assay changes in retinal signaling to increasing amounts of rod loss and accompanying changes in cone morphology and density. Slower forms of RP are also more common among the human population (Grover et al., 1999; Hartong et al., 2006), and thus slow degeneration models may be more therapeutically informative than animal models with relatively rapid photoreceptor degeneration (e.g., *rd1* and *rd10*).

To determine the changes in cone-mediated retinal signaling induced by progressive rod loss, we measured changes in the signaling of retinal ganglion cells (RGCs), the ‘output’ neurons of the retina. Deterioration in the fidelity of RGC signals captures net changes in retinal circuit function induced by rod degeneration: these include changes in retinal circuits that compensate or exacerbate deteriorating photoreceptor function. Large-scale multielectrode arrays (MEAs) were used to measure the visual responses of hundreds of RGCs simultaneously in individual retinas while presenting a variety of visual stimuli: e.g., checkerboard noise and natural movies. Three features of RGC signaling were specifically investigated. First, we measured when and how the spatiotemporal receptive fields (RFs) under cone-mediated conditions were altered by rod degeneration. These RFs are indicative of the visual features that are being signaled by the RGCs to the brain (Chichilnisky, 2001; Keat, Reinagel, Reid, & Meister, 2001; Yu, Grzywacz, Lee, & Field, 2017), and thus changes in these RFs represent changes in the kind of information being transmitted to the brain. Second, we probed how and when the spontaneous activity of RGCs was altered by rod degeneration. Many previous studies have noted the emergence of strong oscillatory activity in the spontaneous activity of RGCs in animal models of RP. Such spontaneous activity could disrupt the ability of the retina to encode stimuli. However, only limited rodent models have been used to identify when these oscillations emerge relative to rod and cone photoreceptor death. These models result from different mutations, which cause distinct changes in rod physiology from those in *Cngb1^neo/neo^* mice; when and how oscillation arise may be mutation specific and may not be ideal for capturing aspects of heterogenous human disease (Stasheff, 2008; Stasheff, Shankar, & Andrews, 2011). Third, we explored when and how rod degeneration impacts the fidelity of visual signals transmitted to the brain. Information theory was used to quantify changes in the fidelity of signaling naturalistic and artificial stimuli as a function of photoreceptor degeneration.

From 1 to 7M of age in *Cngb1^neo/neo^*mice, there were marked and clear changes in cone morphology and density resulting from rod degeneration. When assaying the spatiotemporal RFs of RGCs as a function of degeneration, there were subtle changes to the temporal RFs of RGCs as cone morphology changed. However, the spatiotemporal RFs of RGCs were remarkably stable until the latest stages of degeneration (5-7M). The primary change to cone-mediated RGC responses was a decrease in response gain as photoreceptors were lost. Second, oscillations in the spontaneous activity of RGCs did not emerge in this mouse model until all light responses were eliminated (∼9M). This indicates that the onset of oscillatory activity can depend strongly on the genetic cause of RP, and thus may not be a concern for gene or cell therapy in at least some forms of RP. Third, as rods died and normal cone morphology deteriorated, the ability of the retina to reliably signal the content of natural movies -- under cone-mediated conditions -- was remarkably robust until the latest stages of degeneration (∼7M; total rod loss with severe cone morphology changes and death). Thus, the fidelity of information transmission was relatively stable despite a reduction in response gain. This suggests that homeostatic mechanisms in the retina serve to compensate for deteriorating photoreceptors. Furthermore, these results suggest a broad therapeutic window in which to treat RP, as cone-mediated visual signaling remains surprisingly robust despite clear changes in cone morphology and density.

## Results

### *Cngb1^neo/neo^* cones slowly degenerate lagging rod death

Photoreceptors in *Cngb1^neo/neo^* mice gradually die over a period of 7 to 8M as a consequence of the *Cngb1* mutation (Figure 1A). While rods die first, shown by the shrinking outer nuclear layer (ONL) over time, cones are present until late in disease (Figure 1B). However, cone structure begins to deteriorate at 3M (Figure 1C, middle). Cone outer segments gradually shorten and eventually disappear prior to total cell death (Figure 1D). By 7M, nearly all rods are lost and only cone cell bodies remain; clear outer segment structure is absent (Figure 1C, right). How does this deterioration in cone morphology, and potential changes in cone function impact the ability of the retina to transmit visual information to the brain? Answering this question is particularly important given that humans primarily use cone vision for most tasks. One possibility is that cone-mediated signaling at the level of the RGCs is robust to changes in cone morphology. Alternatively, RGC signaling at cone-mediated light levels may rapidly deteriorate as rods are lost and cone morphology becomes abnormal.

**Figure 1.**
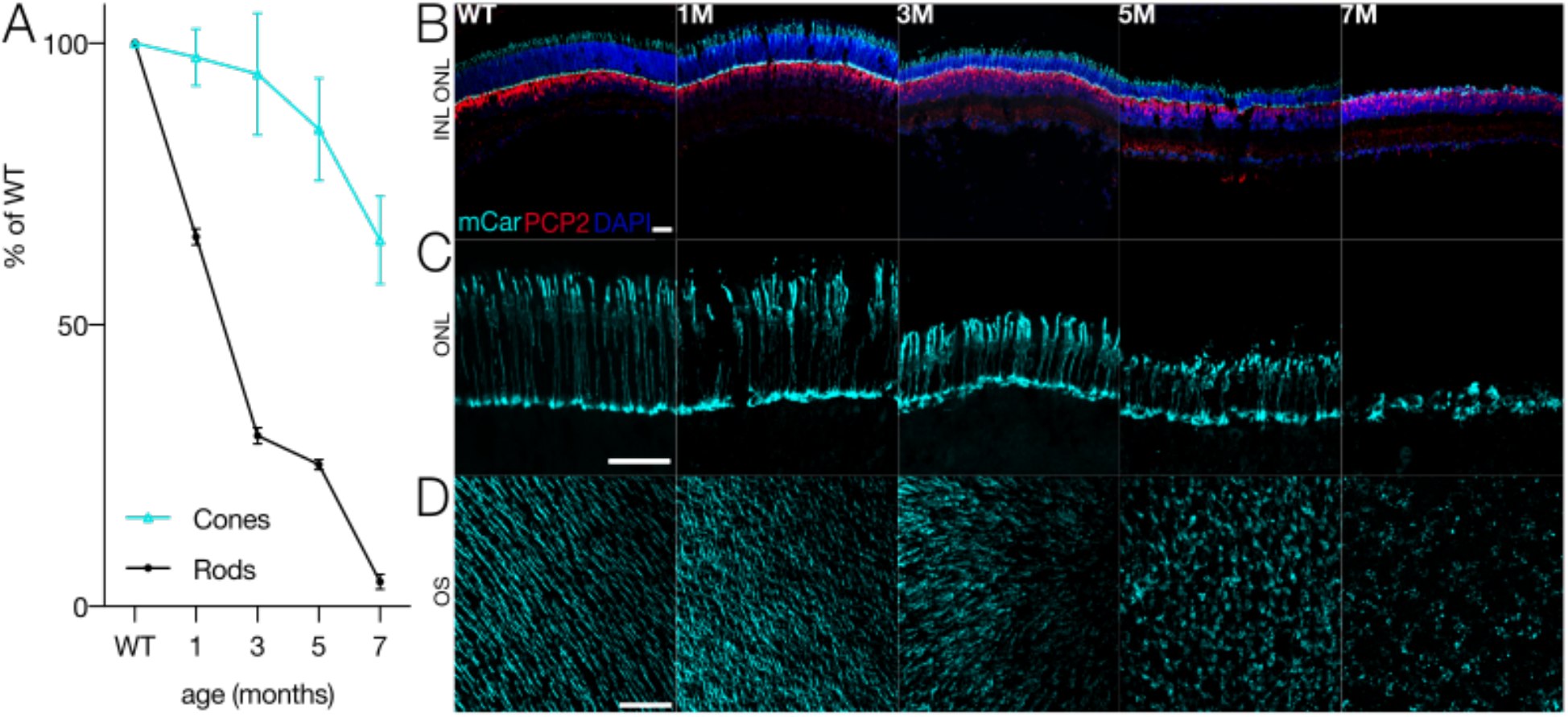
Cone morphology and density change with rod degeneration. A. Estimated fraction of surviving rods (black) and cones (cyan) relative to WT densities from 1 to 7 months of age in *Cngb1^neo/neo^* mice. B. Immunofluorescence of retinal cross sections in WT and *Cngb1^neo/neo^* mice (1-7 months). Cones (mCar) in cyan, ON bipolar cells (PCP2) in red, and nuclei (DAPI) in blue. C. Cone morphologies imaged with cone arrestin (mCar) labeling in WT and *Cngb1^neo/neo^* mice. D. Whole mount view of cone density and morphology in WT and *Cngb1^neo/neo^* mice. Scale bars are 50 µm. ONL: outer nuclear layer; INL: inner nuclear layer; OS: outer segments.

### Identifying RGCs with space-time separable receptive fields

To determine the impact of changing cone morphology and density due to rod death on retinal output, we began by measuring the RFs of RGCs from wild-type (WT) and *Cngb1^neo/neo^* retinas using MEAs consisting of 512 or 519 electrodes (see Methods) (Anishchenko et al., 2010; Field et al., 2010; Litke et al., 2004). The MEAs measured the spiking activity in 186-560 RGCs in individual samples of retina from 31 mice (see Methods). Animals were used at 1M intervals from 1-7M of age (post-natal). To estimate the spatiotemporal RFs of RGCs, we used checkerboard noise and computed the spike-triggered average stimulus (STA) for each RGC (Chichilnisky, 2001).

We measured RF structure at two light levels: a low mesopic light level (100 Rh*/rod/s) at which cones are just beginning to be activated, and a low photopic light level (10,000 Rh*/rod/s). We chose the lower light level because *Cngb1^neo/neo^* mice have severely compromised rod function even in surviving rods, thus we expected to see significant changes in visual signaling between WT and *Cngb1^neo/neo^* (Wang et al., 2019). We chose the higher light level because it largely, if not completely, isolated cone-mediated signaling, thereby allowing us to examine the net impact of rod dysfunction and death on cone-mediated retinal output.

The STA estimates the linear component of each RF. We were particularly interested in changes to the spatial or temporal integration of visual input, which are estimated by changes in the spatial and temporal RFs, respectively. However, some RGC types (i.e., direction-selective RGCs) do not have a RF that can be decomposed into unique spatial or temporal filters (Borghuis, Ratliff, Smith, Sterling, & Balasubramanian, 2008; Devries & Baylor, 1997; Vaney, Sivyer, & Taylor, 2012), which makes it an ill-posed problem to separately analyze changes in spatial or temporal integration. Thus, we focused our analysis on RGCs with RFs that were well-approximated by a space-time separable RF model: the outer product of two vectors, one describing the spatial RF and the other the temporal RF (Figure 2) (see Methods). Singular value decomposition (SVD) factorizes the STA into a linear combination of spatial and temporal filter pairs: the outer product of these pairs, weighted by their associated singular values, will reproduce the original STA (Golomb, Kleinfeld, Reid, Shapley, & Shraiman, 1994; Wolfe & Palmer, 1998). A space-time separable STA will yield one pair of spatial and temporal filters that captures most of the variance in the STA (a.k.a., rank-1 approximation Figure 2A-B). Subsequent space-time filter pairs will exhibit little to no structure and appear as ‘noise’ (Figure 2B, bottom-right). In contrast, non-separable STAs will require multiple space-time vector pairs to reproduce the STA (Figure 2C-D) and each pair will exhibit clear structure in space and time (Figure 2D). RGCs for which > 60% of the variance in the full space-time STA was captured by a separable RF model were included in the analysis. This threshold was chosen because it captured a clear mode in the distribution of all RGCs from WT retinas (Figure 2E). Across time-points of degeneration, 8,028 out of 12,997 RGCs met this criterion. However, the fraction passing this criterion changed as a function of degeneration, with more cells passing the criterion early in degeneration and fewer passing at the latest timepoints (Figure 2F). RGCs passing this criterion had a wide range of spatial RF sizes and temporal RF durations and consisted of both ON and OFF RGC classes. Thus, several, but not all, RGCs met this criterion (see Discussion). Below we analyze changes in the spatial and temporal RFs of RGCs at mesopic and photopic light levels as a function of photoreceptor degeneration.

**Figure 2.**
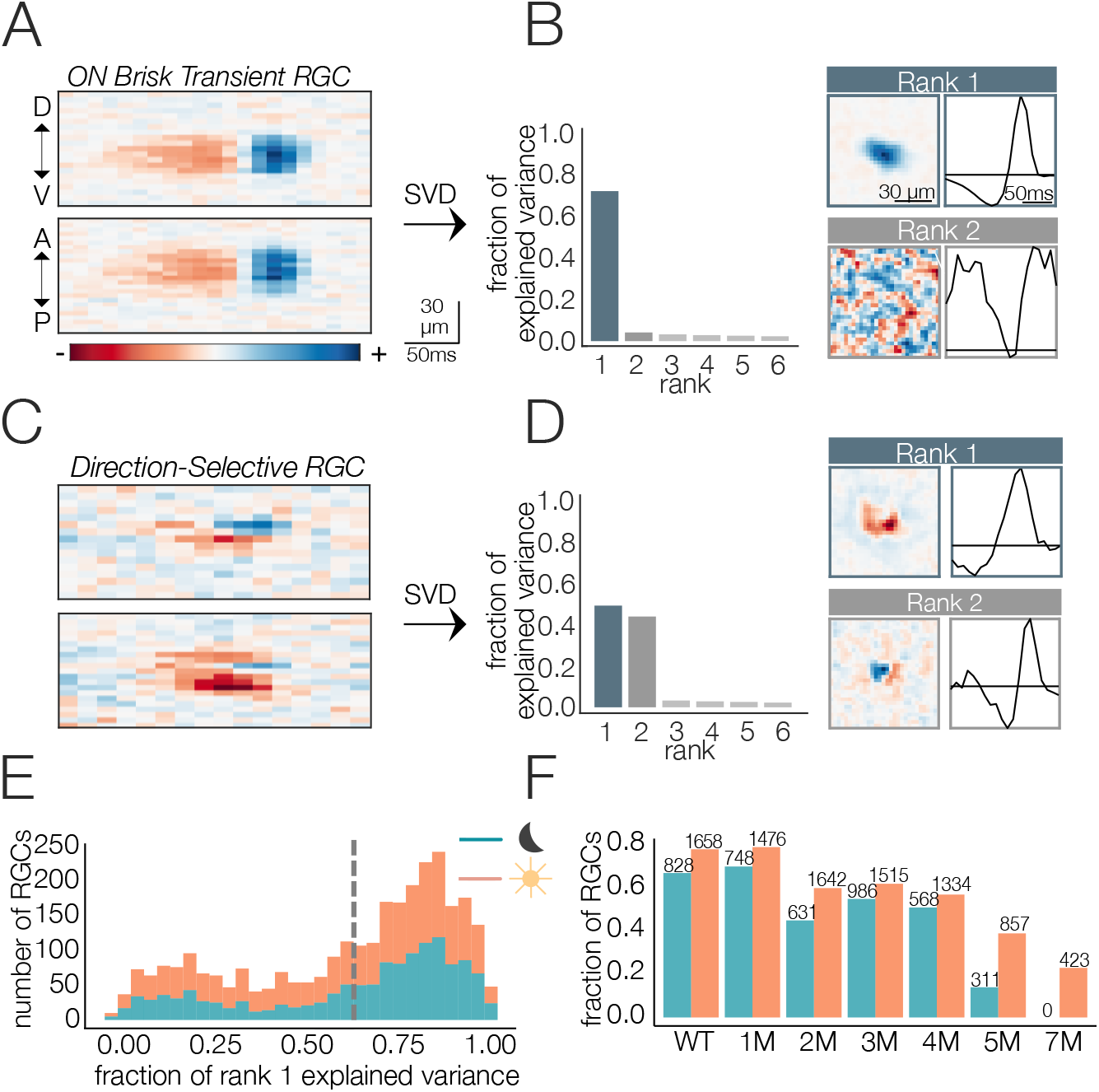
Receptive field decomposition. A. Example space-time separable STA from an RGC. Top is a cut through one spatial dimension and bottom is a cut through the orthogonal spatial dimension. D, V, A, and P indicate the dorsal, ventral, anterior, and posterior directions of the retina. B. Singular value decomposition (SVD) of STA in A: (left) distribution of variance explained by first six space-time vector pairs; (top right: Rank 1) spatial (left) and temporal (right) filters from the rank-one decomposition; (bottom right; Rank 2) spatial and temporal filters associated with the second singular value. C. Example STA from an RGC that was not space-time separable. D. SVD of STA in C; (left) distribution of variance explained by first six space-time vector pairs; (top right; Rank 1) spatial and temporal filters from the rank-one decomposition; (bottom right; Rank 2) spatial and temporal filters associated with the second singular value E. Distribution of variance explained by the rank-one decomposition for all WT RGCs (1654 cells from 5 mice) under mesopic (teal) and photopic (orange) conditions. Vertical line shows threshold for classifying cells as space-time separable. F. Fraction and quantity of RGCs measured at mesopic (teal) and photopic (orange) light levels that were classified as exhibiting space-time separable RFs.

### Changes in mesopic receptive fields with photoreceptor degeneration

At the mesopic light level, the distribution of spatial RF sizes between WT and 1M *Cngb1^neo/neo^* animals was relatively stable (Figure 3A-B). A small, but statistically significant decrease in the mean relative to WT was observed at 1-4M (15% difference between WT to 4M; p-value: 0.039). At 5M, the decrease was attenuated. By 7M, no RGCs exhibited space-time separable RFs at the mesopic light level (Figure 2F) and only 35% of identified RGCs exhibited visual responses (see Methods). Note that corrupting a rank-one STA with increasing amounts of noise will cause the rank-one approximation to capture increasingly less total variance, which likely accounts for the reduced number of RFs at the latest timepoints.

**Figure 3.**
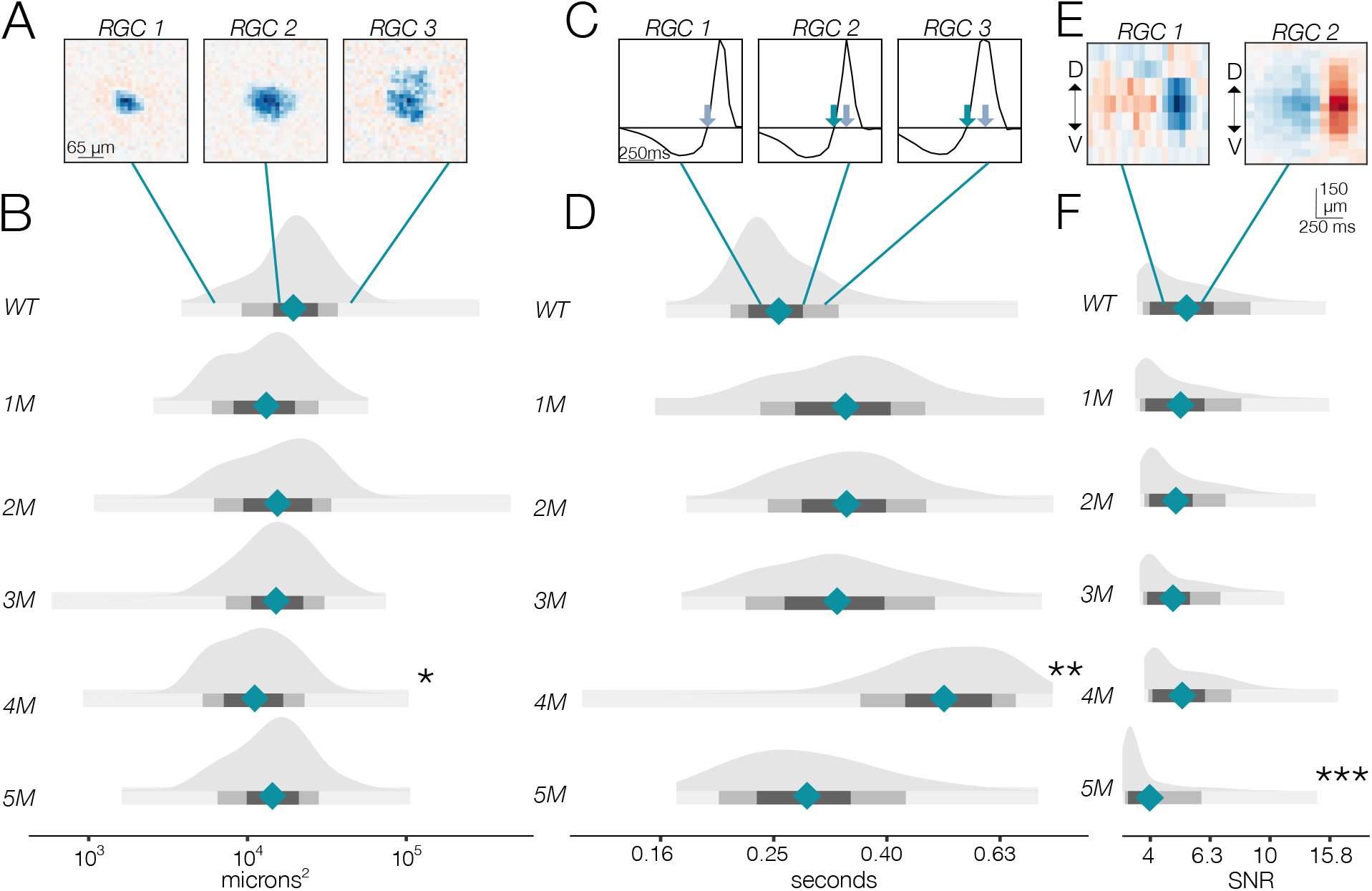
Changes in RF structure under mesopic conditions induced by rod death. A. Example spatial RFs at mesopic light level showing smaller to larger RFs (left to right). B. Distributions of spatial RFs comparing WT and *Cngb1^neo/neo^* mice from 1-5M of degeneration. C. Example temporal RFs at mesopic light level showing shorter to longer temporal integration. Grey arrows indicate the time to zero of RGC 1. Blue arrows on RGC 2 and RGC 3 show time to zero for RGCs 2 & 3. D. Distributions of temporal RF comparing WT and *Cngb1^neo/neo^* mice from 1-5M of degeneration. E. Example STAs with lower (left) and higher (right) signal-to-noise (SNR) ratios. F. Distribution of SNRs comparing WT and *Cngb1^neo/neo^* mice.

At mesopic conditions, we observed larger fractional changes in RGC temporal integration than in their spatial integration between WT and *Cngb1^neo/neo^* mice (Figure 3C-D). At 1-3M the mean temporal integration and the variance was larger for *Cngb1^neo/neo^* animals than WT: mean of all RGCs increased by 14%, 17%, and 13% at 1M, 2M, and 3M respectively (p-value: 0.10, 0.08, 0.11) and the variance of all RGCs increased by 38%, 34%, and 35% at 1, 2, and 3M of age respectively (p-value: 0.01, 0.02, 0.02). At 4M, the median temporal integration increased further by 88% relative to WT (p-value: 0.001). However, at 5M the temporal integration decreased back toward WT (9% increase in time-to-zero relative to WT; p-value: 0.14), with the variance in the distribution substantially larger (37% increase in variance relative to WT; p-value: 0.009). Overall, spatial RF structure was relatively stable until the latest stages of degeneration: the largest change from WT in spatial integration was 15% (at 4M). Changes in temporal integration were greater, with the largest change being 88% (4M), but these changes fluctuated over time and were non-monotonic with degeneration.

Given that photoreceptors are dying, it is potentially surprising that spatiotemporal RF structure is relatively stable under mesopic conditions. Fewer photoreceptors ought to result in diminished sensitivity, even if the area of spatial integration or the duration of temporal integration are relatively stable. Thus, we analyzed the signal-to-noise ratio (SNR) of the STAs which may be expected to be noisier with photoreceptor death. The SNR of the STAs generally drifted down over time (Figure 3E-F); however, even these changes were relatively small until 5M (compared to WT there was a 5% and 15% decline in SNR in the 4M and 5M experimental groups, respectively; p-value: 0.19 and 0.04). Of note, the STA measurements were based on 30 min of checkerboard stimuli, which may be a sufficiently long period of time to average away noise and/or compensate for diminished gain. To more directly assess changes in response gain, we inspected the contrast response functions (also called ‘static nonlinearities’) across RGCs, which capture the relationship between the stimulus (filtered by the RF) and the spiking output of the RGC (Chichilnisky, 2001). The gain of these contrast response functions steadily decreased as a function of degeneration (Figure 4). The effect was most noticeable for stimuli that were the most similar to the RF (Figure 4C-E). Thus, under mesopic conditions, gain steadily decreased with photoreceptor degeneration, while spatial and temporal integration were relatively stable until the latest stages of degeneration.

**Figure 4.**
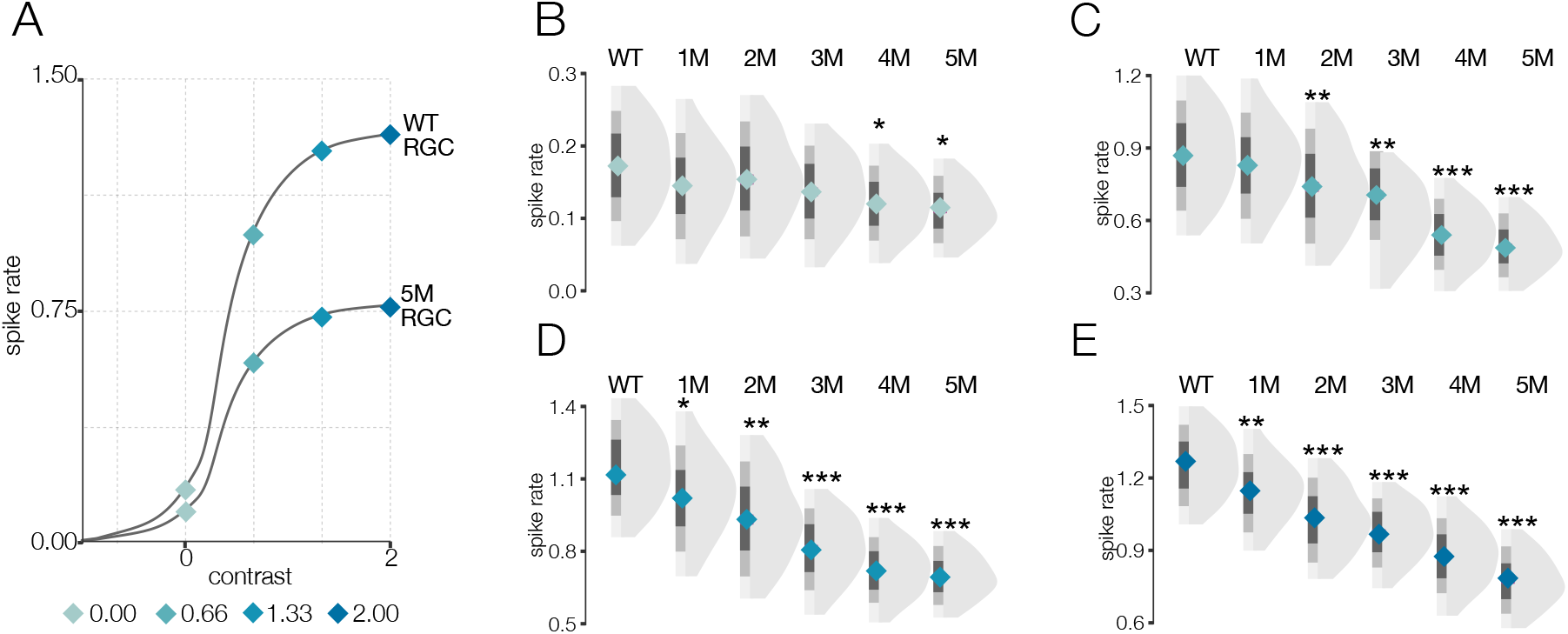
Gain of contrast response functions steadily decreases during RP under mesopic conditions. A. Mean cumulative Gaussian fit to contrast response function across RGCs from WT and 5M *Cngb1^neo/neo^* mice. Four locations along the contrast response functions are highlighted by blue diamonds. B-E, Distribution of contrast response function values at contrast values indicated in A for WT and 1-5M retinas. Note, this analysis was not restricted to RGCs with space-time separable RFs, allowing the analysis to extend to 5M.

### Changes in photopic receptive fields with photoreceptor degeneration

Under photopic conditions, the distribution of spatial RF sizes was remarkably stable throughout degeneration (Figure 5A-B). The largest change was at 7M with a 5% decrease in spatial RF area relative to WT (p-value: 0.211). Note, unlike the mesopic condition, RFs were detectable out to 7M under the photopic condition.

**Figure 5.**
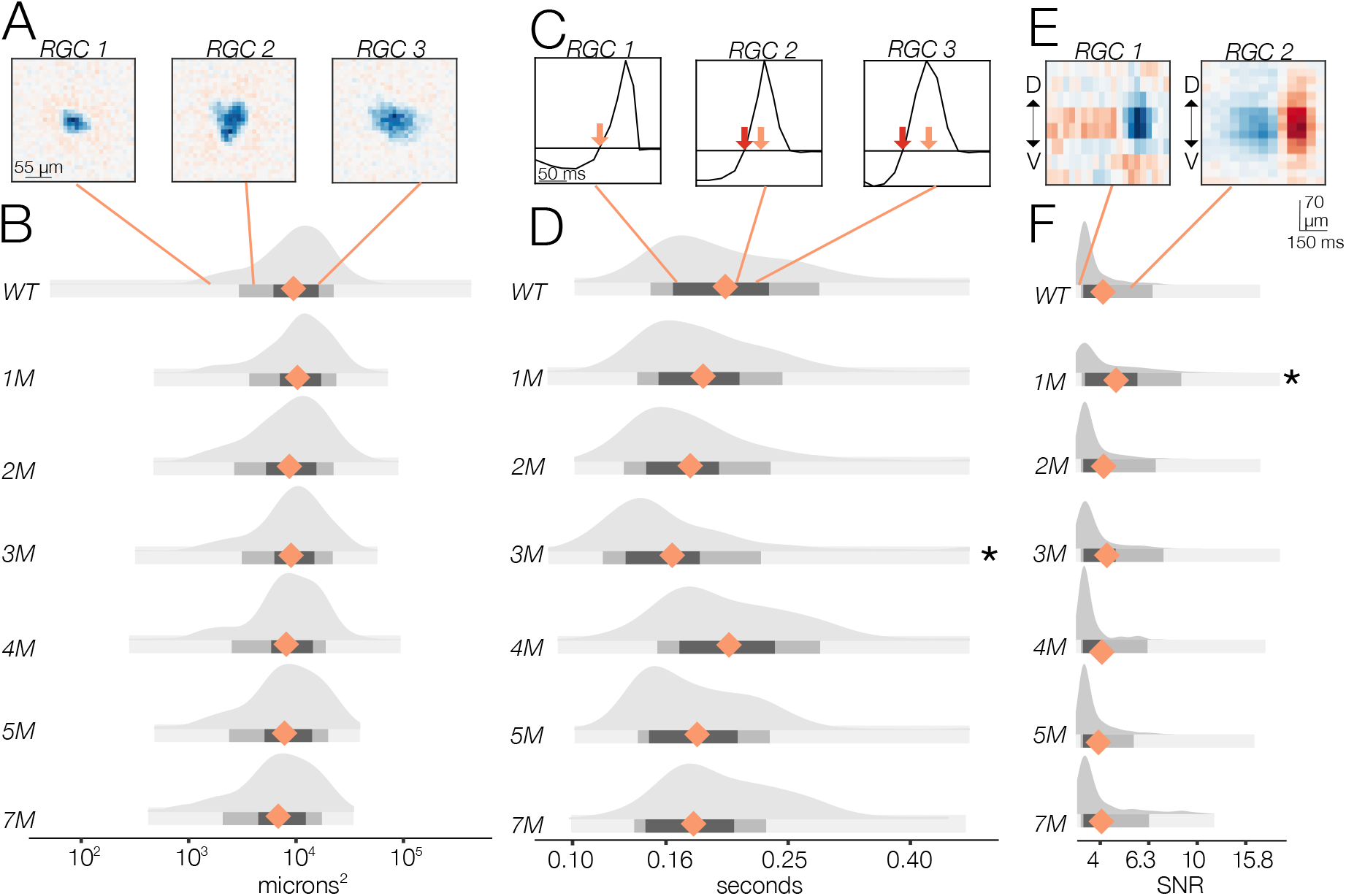
Changes in RF structure under photopic conditions induced by rod death. A. Example spatial RFs at the photopic light level showing small to larger RFs (left to right). B. Distributions of spatial RFs comparing WT and *Cngb1^neo/neo^* mice from 1-7M of degeneration. C. Example temporal RFs at photopic light level. Orange arrows indicates the time to zero for RGC 1; Red arrows show time to zero for RGCs 2 and 3. D. Distributions of temporal RFs comparing WT and *Cngb1^neo/neo^* mice from 1-7M of degeneration. E. Example STAs with lower and higher signal-to-noise (SNR) ratios. F. Distribution of SNRs comparing WT and *Cngb1^neo/neo^* mice.

Temporal RFs exhibited greater changes than spatial RFs under photopic conditions (Figure 5C-D) but were still surprisingly stable. Specifically, the time to zero progressively decreased 10%, 12%, and 15% from 1-3M, respectively (relative to WT; p-value: 0.32, 0.22, 0.04), indicating a shortening of temporal integration. At 4M the temporal integration slowed somewhat, becoming 22% slower than 3M but only 4% slower than WT (p-value: 0.423) (Figure 5D). At 5M and 7M, time to zero decreased by 12% and 13% (p-value: 0.22, 0.12), respectively, relative to 4M.

These changes in temporal and spatial RF structure under photopic conditions were relatively small given the clear changes in cone morphology observed particularly at 5 to 7 M (Figure 1): the largest deviation in mean temporal integration from WT occurred at 3M, and this was only a 15% slowing in the time-to-zero, while spatial RF sizes changed by 5% at most (relative to WT) through 7M.

We next analyzed the SNR of the STAs (Figure 5E-F). We observed a decline in SNR with degeneration under photopic conditions. Yet even at 7M, with 30% cone loss, near complete rod loss, and abnormal cone morphologies, there was only a 6% decrease in the median SNR (Figure 5F; p-value: 0.456). As with the mesopic conditions, computing STAs over relatively long periods of checkerboard noise may mask decreases in gain or increases in response noise. To inspect changes in gain, we analyzed the contrast response functions (Figure 6). Like the mesopic condition, response gain steadily decreased with degeneration, particularly for stimuli that increasingly matched the RF (Figure 6D-E). Thus, the dominant effect of photoreceptor degeneration on photopic RF structure among RGCs in *Cngb1^neo/neo^* mice was a decrease in response gain, not a change in spatial or temporal integration.

**Figure 6.**
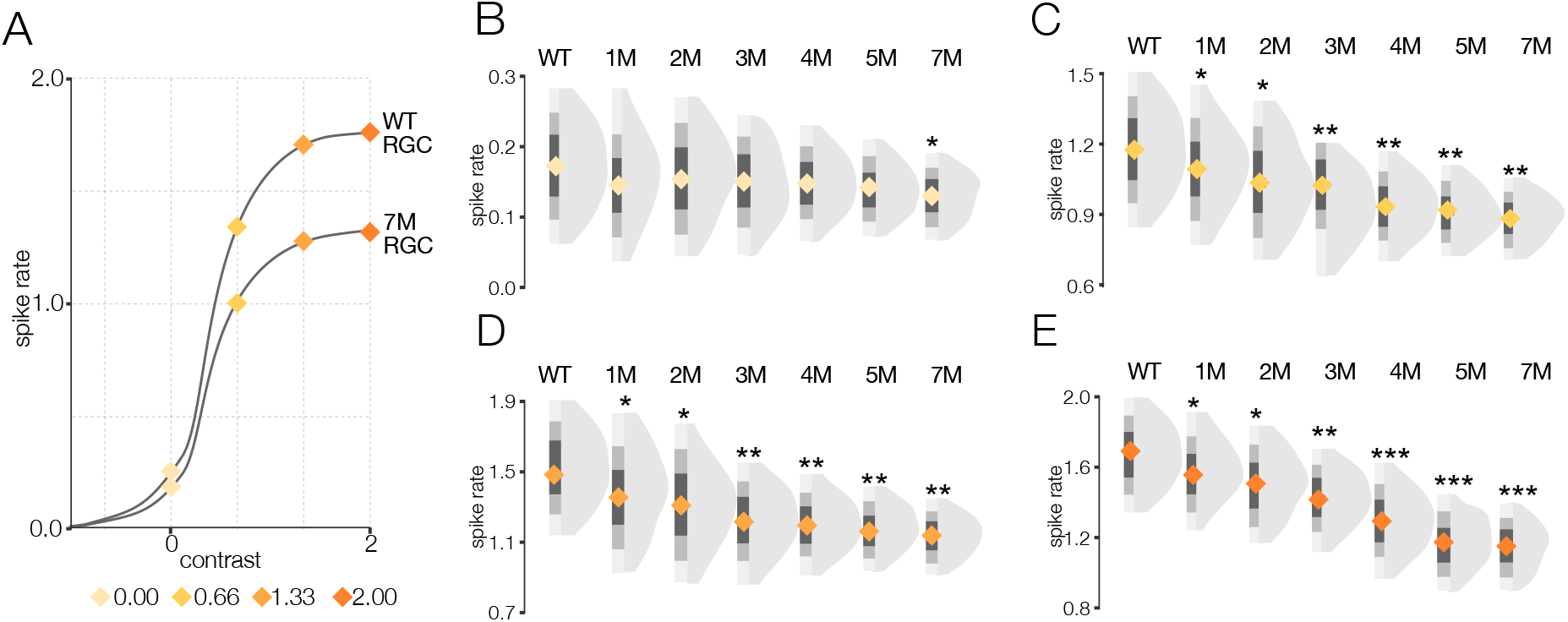
Gain of contrast response functions steadily decreases during RP under photopic conditions. A. Mean cumulative Gaussian fit to contrast response function across RGCs from WT and 7M *Cngb1^neo/neo^* mice. Four locations along the contrast response functions are highlighted by diamonds. B-E, Distribution of contrast response function values at contrast values indicated in A for WT and 1-7M retinas.

### *Cngb1^neo/neo^* RGCs do not exhibit oscillatory activity while photoreceptors remain

The results above indicate relatively small changes in spatial and temporal integration of cone mediated responses as rods die and cones degenerate in *Cngb1^neo/neo^*mice. They also indicate clear decreases in gain as a function of degeneration under mesopic and photopic conditions (Figures 4 and 6). However, the analyses above do not provide much insight into the extent that noise may be changing with degeneration. One source of noise is signal-independent increases in spontaneous activity. Several rodent models of retinal degeneration exhibit increased spontaneous activity arising as 5-10 Hz oscillations in RGC spiking (Marc et al., 2007; Margolis, Newkirk, Euler, & Detwiler, 2008; Stasheff, 2008). These oscillations are reported to occur throughout the degeneration process (Stasheff et al., 2011). The presence of such oscillations may minimally impact the RF measurements as it would be largely averaged away when computing the STA, but they could contribute to the lower SNR of the STAs (Figure 3F and 5F) along with the decreased gain (Figures 4 and 6).

Thus, we analyzed the spontaneous activity of RGCs in WT and *Cngb1^neo/neo^* animals from 1-9M to check for oscillatory activity (Figure 7). Spontaneous activity was measured in darkness as well as with static illumination set to the same mean intensity as the checkerboard stimuli presented at the mesopic and photopic light levels. Oscillations were absent in the spiking activity of all recorded RGCs in darkness and at both mesopic and photopic light levels in *Cngb1^neo/neo^* retinas from 1 – 7M (Figure 7A-D, G). However, clear 5 Hz oscillatory activity did arise at 9M (Figure 7E-G). The oscillations were present at all three light levels (not shown). At this last stage of degeneration, no light response could be detected with 100% contrast flashes at the mesopic or photopic light levels (not shown). Thus, oscillations in the spontaneous spiking activity of RGCs in the *Cngb1^neo/neo^* mouse model are absent until all (or nearly-all) of the RGCs have lost their light responses.

**Figure 7.**
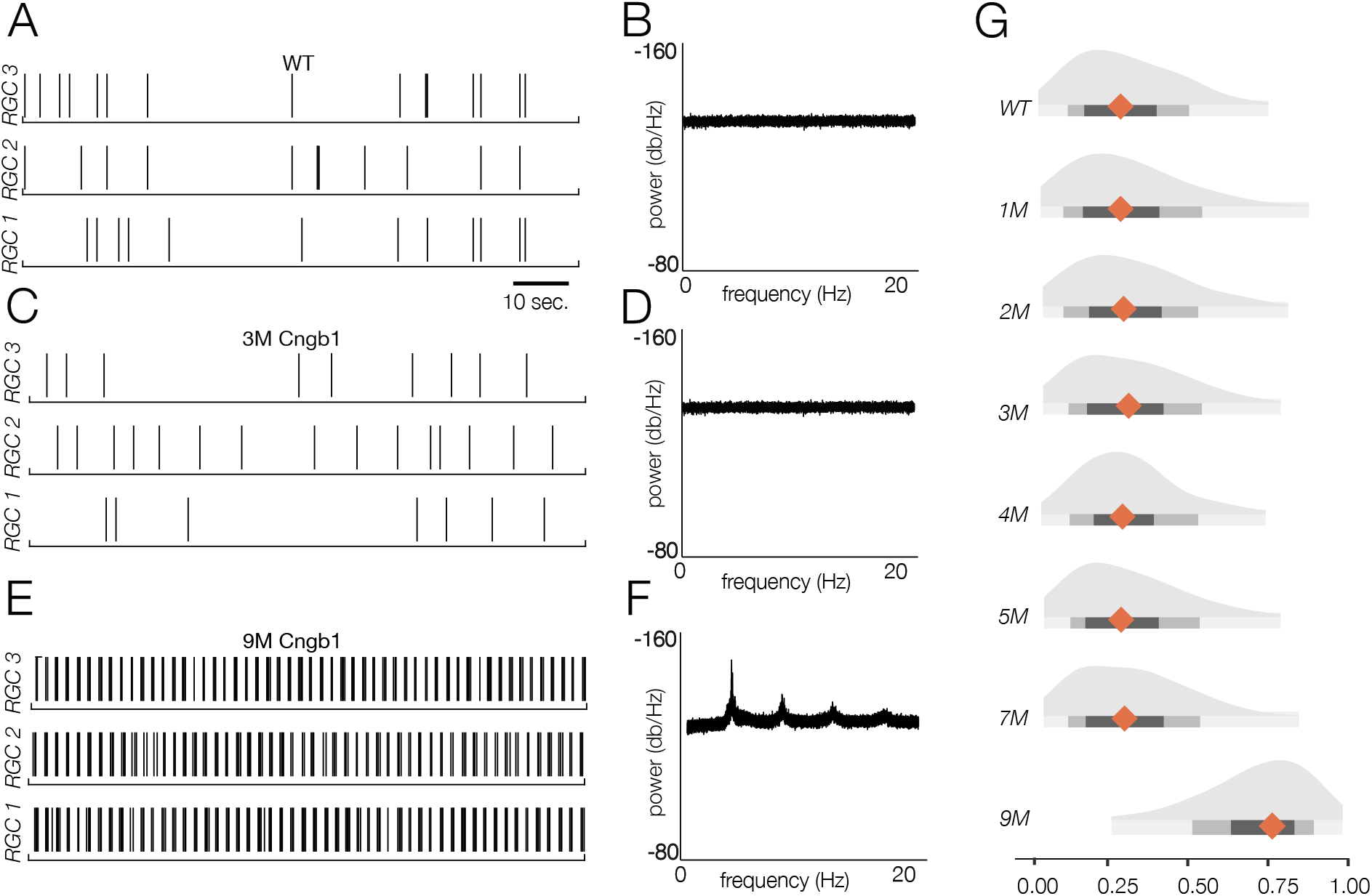
Retinal oscillations occur after vision loss in *Cngb1^neo/neo^* mice. A. Spontaneous activity of three representative WT RGCs in total darkness. B. The power spectral density (PSD) of one example RGC. C-D. Example rasters (C) and PSD (D) of a representative 3M *Cngb1^neo/neo^* RGC. E-F. Example rasters (left) and the PSD of an example 9M *Cngb1^neo/neo^* RGC. G. Distributions of the PSD fluctuation ratio, defined as the maximum of PSD divided by the baseline PSD (see Methods). Mean indicated by orange diamond, gray bars indicate 100%, 80%, and 50% sections of the distribution around the median.

### RGCs from *Cngb1^neo/neo^* retinas exhibit deteriorated visual signaling under mesopic conditions

Thus far, we have observed a decrease in response gain among RGCs under mesopic and photopic conditions that tracks progressive photoreceptor degeneration. However, signal-independent noise, as assayed by changes in spontaneous activity, appears constant until all (or nearly all) photoreceptors have died in *Cngb1^neo/neo^* mice. It is possible that the RGCs are signaling less reliably as photoreceptors die: the STA method averages over many trials and is thus relatively insensitive to potential changes in signal-dependent noise that could manifest as increased variability in either the number or timing of spikes produced to a given stimulus. Because of this, it is important to identify potential changes in the fidelity of RGC signaling that may accompany photoreceptor degeneration.

To measure the fidelity of RGC signaling, we used information theory (Shannon, 1948). Specifically, we calculated the mutual information between spike train and the stimulus (see Methods). Mutual information indicates how much the uncertainty about the visual stimulus is reduced by observing the spike train of an RGC (see review by McDonnell, Ikeda, & Manton, 2011). If the RGC response is highly variable, observing a single response will reduce uncertainty about the stimulus less than for an RGC with highly reproducible responses.

We began by presenting a repeating checkerboard stimulus at the mesopic light level used for the RF measurements above (∼100 Rh*/ rod/ s). We segregated RGCs based on their information rate for the checkerboard stimulus (Figure 8). Some RGCs exhibited strongly modulated and reproducible responses across repetitions of the stimulus (Figure 8A, RGC 1), while other RGCs were less reliably driven by the stimulus (Figure 8A, RGC 2 & 3). Note all RGCs with a spike rate above 3 Hz (12,997 RGCs) were used in this analysis: we included RGCs with and without space-time separable RFs.

**Figure 8.**
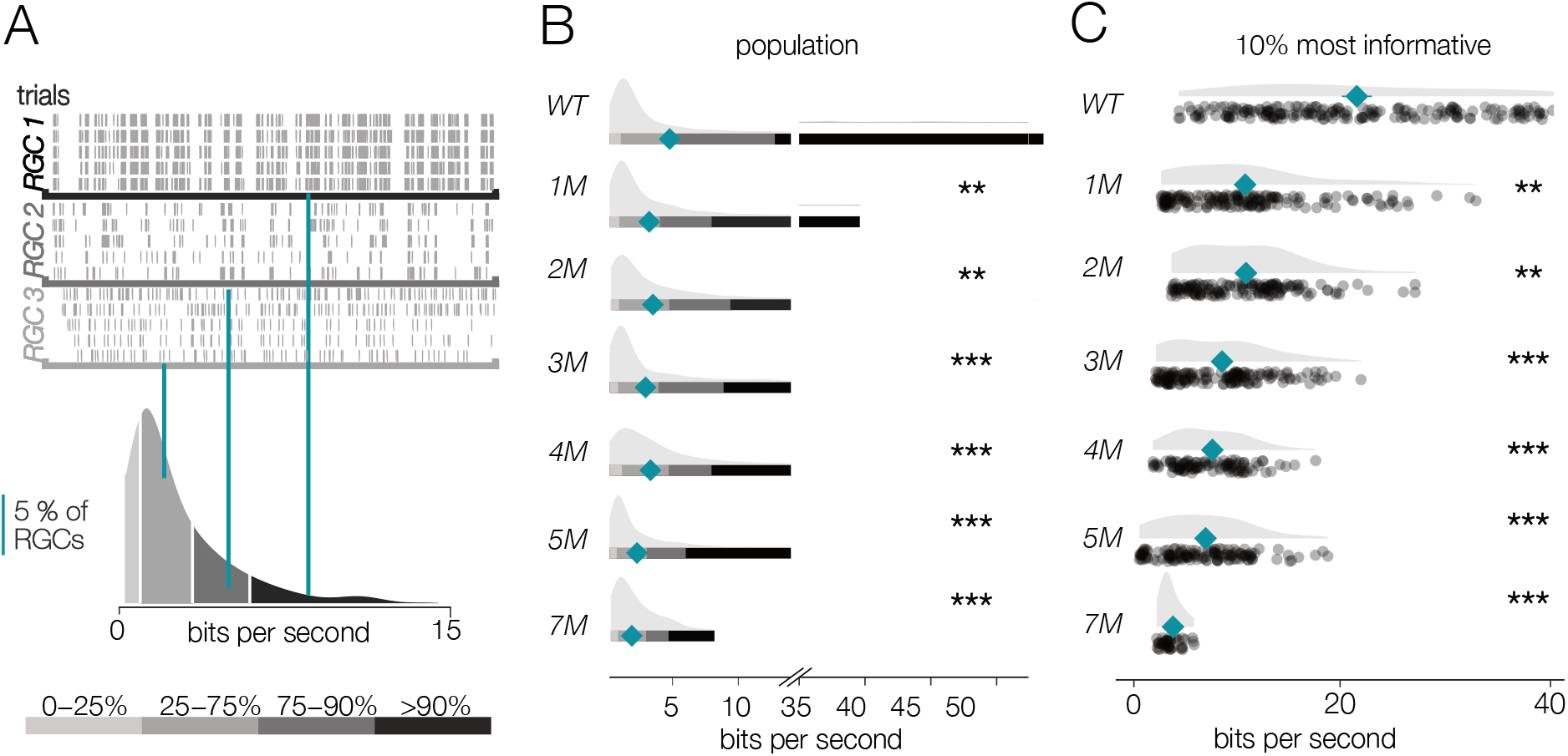
RGC signaling fidelity under mesopic conditions decreases with photoreceptor degeneration. A. (Top) Rasters from three example RGC responding to a repeated mesopic checkerboard stimulus. (Bottom) distribution of information rates across all RGCs from 1 experiment; teal lines show where each example RGC lies in the distribution. B. Distributions of information rates of for all RGCs in each condition: WT and *Cngb1^neo/neo^* mice. Mean shown by teal diamond, stars indicate significant changes from WT: ** and *** are p < 0.01 and 0.001, respectively. C. Distribution of information rates of 10% most informative RGCs across conditions: WT and *Cngb1^neo/neo^* mice.

For WT retinas, the distribution of information rates across RGCs was approximately unimodal with a long tail skewed toward high rates (Figure 8A, bottom). The peak of these distributions did not change much with degeneration (Figure 8B). However, the peak primarily represented cells that were not particularly reliably driven by the checkerboard stimulus even in WT retinas; in other words, these were not the cells that were most informative about the stimulus. In contrast, the median of the information rate distributions decreased as photoreceptors died (Figure 8B, teal diamonds). This change in the median was largely driven by decreased information rates in the long tails of these distributions (Figure 8B). We thus focused our analysis on the 10% of RGCs that were most informative about the checkerboard stimulus: those with the highest information rates (Figure 8C). For these RGCs, there was a clear drop in information rates between WT and 1M *Cngb1^neo/neo^* mice (mean of all RGCs changed from 5.7 bits/s to 4.3 bits/s; p-value: 0.003). In fact, *Cngb1^neo/neo^* RGCs at all time points examined transmitted significantly less information than WT RGCs at the mesopic light level (mean of all RGCs ranged from 2.1-4.3 bits/s). Information rates between 1 and 2M did not change significantly (difference of 0.09 bits/second was observed; p-value: 0.655), but there was a steady decline at subsequent timepoints.

These data indicate that under low mesopic conditions, there is a relatively steady decline in the reliability with which RGCs signal visual information in *Cngb1^neo/neo^* mice. The difference between WT and 1-2M *Cngb1^neo/neo^* animals is likely because rods are poorly responsive to light in *Cngb1^neo/neo^*animals due to a greatly reduced photocurrent (Wang et al., 2019). The progressive decline in information rates after 2M is likely caused by large-scale loss of rods (Figure 1A), and possibly a reduced ability of cones to signal near their threshold for activation.

### RGCs from *Cngb1^neo/neo^* retinas exhibit relatively stable visual signaling under photopic conditions

We next checked the reliability of RGC signaling under photopic conditions that are well above the threshold of cone vision (10,000 R*/rod/s). It is possible that at higher light levels, visual signaling is more (or less) stable. Similar to the lower light level, RGCs exhibited a wide range of information rates to checkerboard stimuli, with some exhibiting very reliable responses to a repeated stimulus while others exhibited more variable responses (Figure 9A). Information rates were much more stable among RGCs at these higher light levels, with no significant differences in median information rates observed between WT and 1M to 3Ms of degeneration (p-values of 0.56, 0.32, 0.29 between WT and 1M, 2M, and 3M, respectively) (Figure 9B). Even when focusing on the top 10% of most informative RGCs, 1 and 2M of degeneration were not significantly different from WT (Figure 9C). Despite clear changes in cone morphology at 5Ms (Figure 1), decreases in information rates were modest (mean of top 10% of RGCs changed from 23.1 bits/second in WT to 18.1 bits/s at 5M; p-value: 0.0009). At 7M, cone-mediated signaling among the RGCs continued to transmit information, but with a relatively pronounced decrease: mean rate was 8.6 bits/s among the top 10% of RGCs. Overall, these data indicate that the fidelity of cone-mediated visual signaling among RGCs is quite stable until ∼2-3M of degeneration, corresponding to 50-70% rod loss.

**Figure 9.**
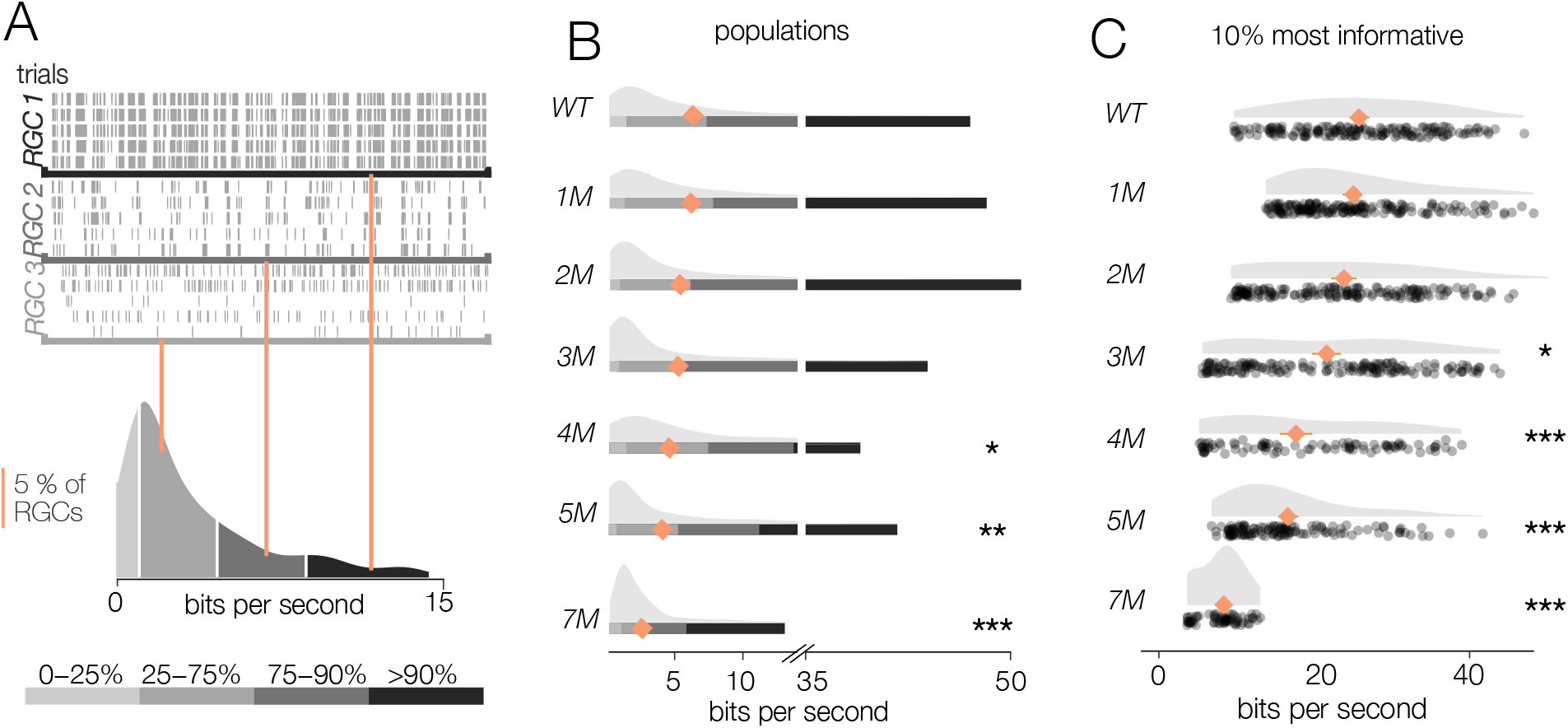
RGC signaling fidelity under photopic conditions is relatively stable with rod degeneration. A. (Top) Rasters from three example RGC responding to a repeated photopic checkerboard stimulus. (Bottom) distribution of information rates across all RGCs from 1 experiment; teal lines show where each example RGC lies in the distribution. B. Distributions of information rates of for all RGCs in each condition: WT and *Cngb1^neo/neo^* mice. Mean shown by teal diamond, stars designate significant changes from WT: ** and *** are p < 0.01 and 0.001, respectively. C. Distribution of information rates of 10% most informative RGCs across conditions: WT and *Cngb1^neo/neo^* mice.

### Degenerating retina more robustly encodes natural than artificial stimuli

Given the wide range of information rates observed across RGCs for checkerboard stimuli, we hypothesized that the results described above may depend on stimulus choice. Thus, we repeated the experiments under photopic conditions (10,000 Rh*/rod/s) using 2 natural movies (see Methods). As with the checkerboard stimuli, RGCs again demonstrated a wide range of information rates when presented with the natural movies (Figure 10A). However, for natural movies, the median information rates were much more stable as photoreceptor degeneration progressed (Figure 10B): relative to WT, there was only a 5% decline in MI at 4M; p-value: 0.543). Similarly, the RGCs with the highest information rates (top 10%) did not show the same decline in information rate as observed with the checkboard stimuli (Figure 10C): from WT to 4M, there was only a 9.5% decline in information rate (p-value: 0.321).

**Figure 10.**
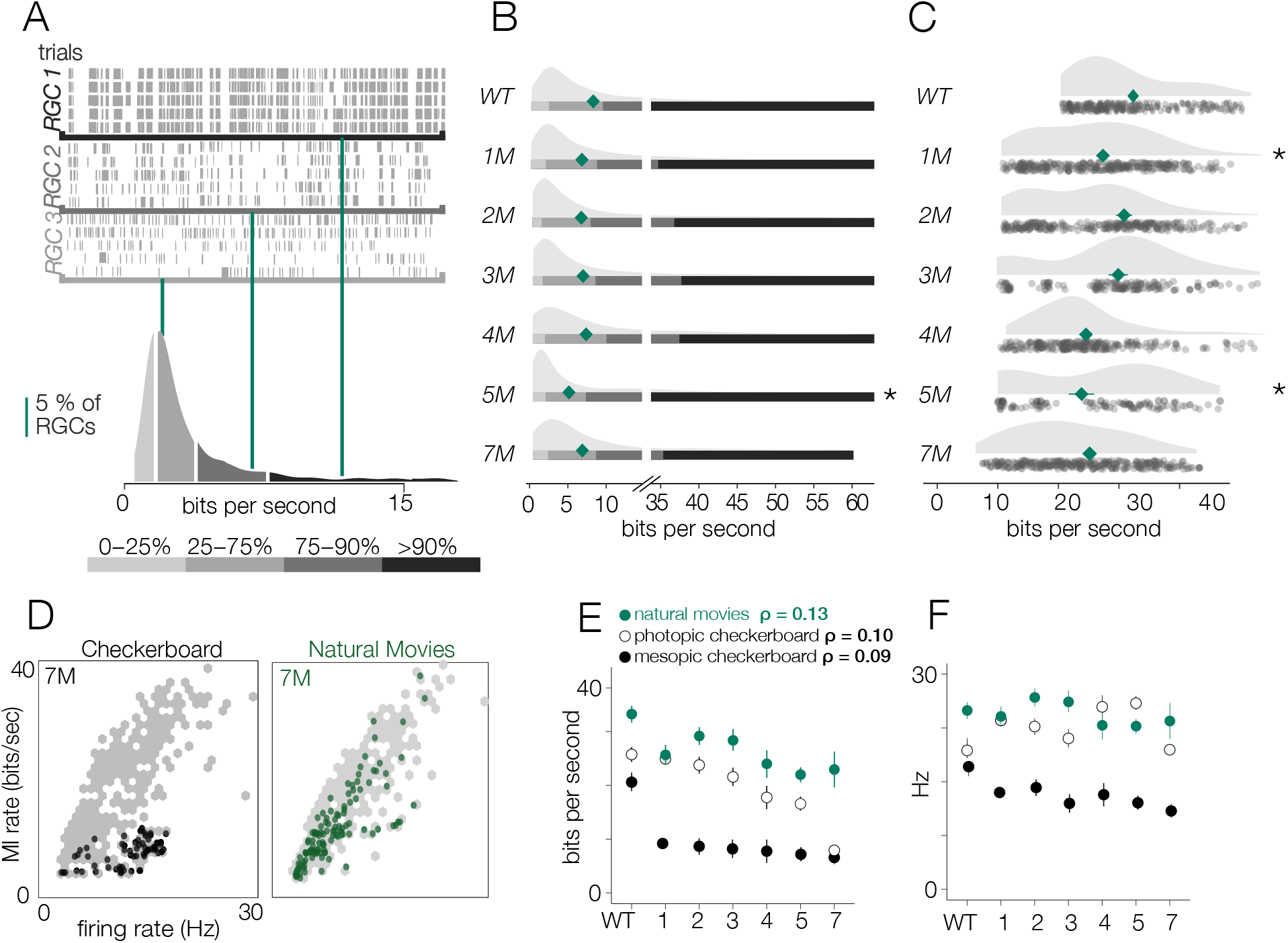
*Cngb1^neo/neo^* RGCs signal more information about natural movies than checkerboard movies. A. (Top) Rasters from three example RGCs responding to a repeated photopic natural stimulus. (Bottom) distribution of information rates across all RGCs from 1 experiment; green lines show where each example RGC lies in the distribution. B. Distributions of information rates of for all RGCs in each condition for WT and *Cngb1^neo/neo^* mice. Mean shown by green diamond. C. Distribution of information rates of 10% most informative RGCs across conditions: WT and *Cngb1^neo/neo^* mice. D. Scatter plot of information rates and spike rates from RGC responses to photopic checkerboard stimulus (left) or natural stimuli (right). Grey dots are RGC responses from the total population, while black dots are specific to 7M *Cngb1^neo/neo^* mutual responses to a checkerboard stimuli and green dots 7M *Cngb1^neo/neo^* responses to a natural movie. E. (left) Mean +/- 2 standard error of information rates of RGC responses to natural movies (green), photopic checkerboard (open), and mesopic checkerboard (black) across WT and *Cngb1^neo/neo^* mice. (right) Mean +/- 2 standard error of spike rates of RGC responses to natural movies (green), photopic checkerboard movies (open), and mesopic checkerboard (black) across WT and *Cngb1^neo/neo^* mice. ρ is the linear correlation between the mean MI and the mean firing rate across retinas.

Finally, we inspected how the information rates depended on the spike rates. Lower information rates could be caused by stimuli producing fewer spikes in degenerating retinas, and stimuli induced fewer spikes at later stages of degeneration (Figure 6). Thus, we compared the information rate to the firing rate for checkerboard stimuli and the natural movies (Figure 10D). When comparing information rates across all RGCs at all timepoints, there is a clear correlation between spike rate and information rate for the checkerboard stimuli and natural movies (Figure 10D, gray dots). However, at 7M for checkerboard stimuli (Figure 10D, black dots), RGCs with the highest firing rates exhibited suppressed information rates relative to RGCs with similar firing rates from retinas with less degeneration. For natural movies, RGCs from 7M retinas exhibited much higher information rates across a similar range of firing rates observed in checkerboard stimuli (Figure 10D). Inspecting across degeneration timepoints, there was a sharp drop in the information rates for mesopic checkerboard stimuli from WT to 1M *Cngb1^neo/neo^*animals (10% most informative cells) that was not present under photopic conditions (Figure 10E). Under photopic conditions, information transmission was higher among the 10% most informative cells for natural movies than for checkerboard stimuli (Figure 10E). These changes in information rates were not strongly correlated with the changes in spike rates across conditions (Figure 10F, ρ = 0.13), suggesting that they are not simply a result of changing spike rates.

## Discussion

Determining the extent to which rod dysfunction, degeneration, and death impacts cone-mediated visual signaling in the retina is important for diagnosing and treating RP. It is possible that cone-mediated signals in RGCs, the ‘output’ neurons of the retina, rapidly deteriorate with the loss of rods and concomitant changes in cone morphology. Alternatively, it is conceivable that RGC signaling is robust to photoreceptor degeneration. We have measured cone-mediated visual responses from thousands of RGCs from *Cngb1^neo/neo^* mice as rod and cone photoreceptors degenerate. We have spanned early timepoints at which most rods remain and there is little to no cone death or changes in cone morphology, to time points at which all rods are lost and cones exhibit abnormal morphologies and are beginning to die. We find that despite clear changes in cone morphology and density, RF structure and signal fidelity remain relatively stable until the latest stages of degeneration. In this mouse line, oscillatory spontaneous activity among RGCs did not arise until all or nearly all photoreceptors were lost in contrast to previous results from *other RP models*. These results suggest that early rescue of rods from degeneration is likely to preserve relatively normal cone-mediated visual signaling among RGCs.

### Comparison to previous studies of RGC signaling in retinitis pigmentosa

Few studies track cone-mediated RGC signaling at many timepoints across RP. Focusing on RGC signaling has the advantage that it includes degradation in visual signaling caused by the degeneration of photoreceptors as well as downstream changes in retinal circuitry. Thus, it captures net changes in retinal processing that deteriorate signaling along with homeostatic mechanisms that may serve to preserve signaling (Care et al., 2020, 2019; Leinonen et al., 2020; Shen et al., 2020). In addition, examining cone-mediated vision, instead of rod vision, focuses on changes that are most likely to be relevant for humans. We did not distinguish among RGC types, other than identifying those with space-time separable RFs for the spatial and temporal RF analyses (Figures 2-6). This choice was largely because at 5M-7M it became increasingly difficult to reliably distinguish cell types, and at the earlier time points we did not observe systematic differences across RGCs types.

Previous work in a *P23H* rat model of RP indicated a monotonic decrease in RF size, a monotonic increase in the duration of temporal integration, and a rise in spontaneous activity as photoreceptors died (Sekirnjak et al., 2011). In contrast, using *Cngb1^neo/neo^* mice we observed small shifts in RF size and temporal duration during degeneration, but these changes were not monotonic with disease progression. The only feature of visual signaling that we found to change monotonically was response gain (Figures 4 & 6). Both studies found that cone-mediated responses persisted longer than rod-mediated responses, which is unsurprising given that cones typically survive longer than rods in RP. We speculate that the differences between these studies arise from degeneration having different causes in *P23H* rats versus *Cngb1^neo/neo^* mice. The rat model involves a mutation in rhodopsin which causes issues with protein folding and trafficking that are not thought to initially change the dark-current of rods or their resting glutamate release (Jones & Marc, 2005; Liu, Garriga, & Khorana, 1996; Machida et al., 2000; Sakami et al., 2011). However, the *Cngb1^neo/neo^* rods exhibit a reduced dark current, are tonically hyperpolarized, and likely release less glutamate onto post-synaptic bipolar cells (Wang et al., 2019). These differences in rod physiology and how rods signal to downstream circuits are likely to have consequences for how the retina responds to rod death. Fully understanding how different origins of rod degeneration impact retinal circuits and RGC signaling is an important direction for future RP studies.

A novel feature of this study is the use of information theory to assay the reliability of visual signaling during retinal degeneration (Figures 8-10). Information theory has been used extensively in the vertebrate and invertebrate visual systems, as well as other sensory systems to quantify encoding performance of sensory neurons (Fairhall et al., 2006; K. Koch et al., 2006; Rieke, Bodnar, & Bialek, 1995; Van Steveninck, Lewen, Strong, Koberle, & Bialek, 1997). In general, sensory neurons with more reliable and selective responses exhibit higher information rates, measured in bits per second. Information theory has not been used previously to assay the effects of neural degeneration on sensory coding. Information theory has the advantage in that it provides a way of comparing the signal fidelity across neurons that transmit information about very different sorts of features (e.g., color vs. motion). It has the potential disadvantages that it is a data-hungry analysis and can be prone to certain biases (Paninski, 2003). The relatively long and stable nature of MEA experiments allowed us to overcome these limitations (Figure 11). Consistent with previous results, we found a broad range of information rates across the population of RGCs (K. Koch et al., 2004, 2006), even in control (WT) retinas. What was perhaps surprising is how stable these information rates were under photopic conditions until those latest stages of degeneration. This indicates that while gain diminishes with degeneration (Figures 4 & 6), the precision and reliability of RGC spiking remains relatively stable (Figures 8-10). Interestingly, this reliability of signaling was more robust when using natural stimuli compared to checkerboard noise (Figure 10). This difference may be the result of natural stimuli having spatiotemporal correlations that are not present in checkerboard noise, and homeostatic mechanisms that preserve retinal signaling during degeneration can potentially exploit these correlations for higher fidelity signaling.

**Figure 11.**
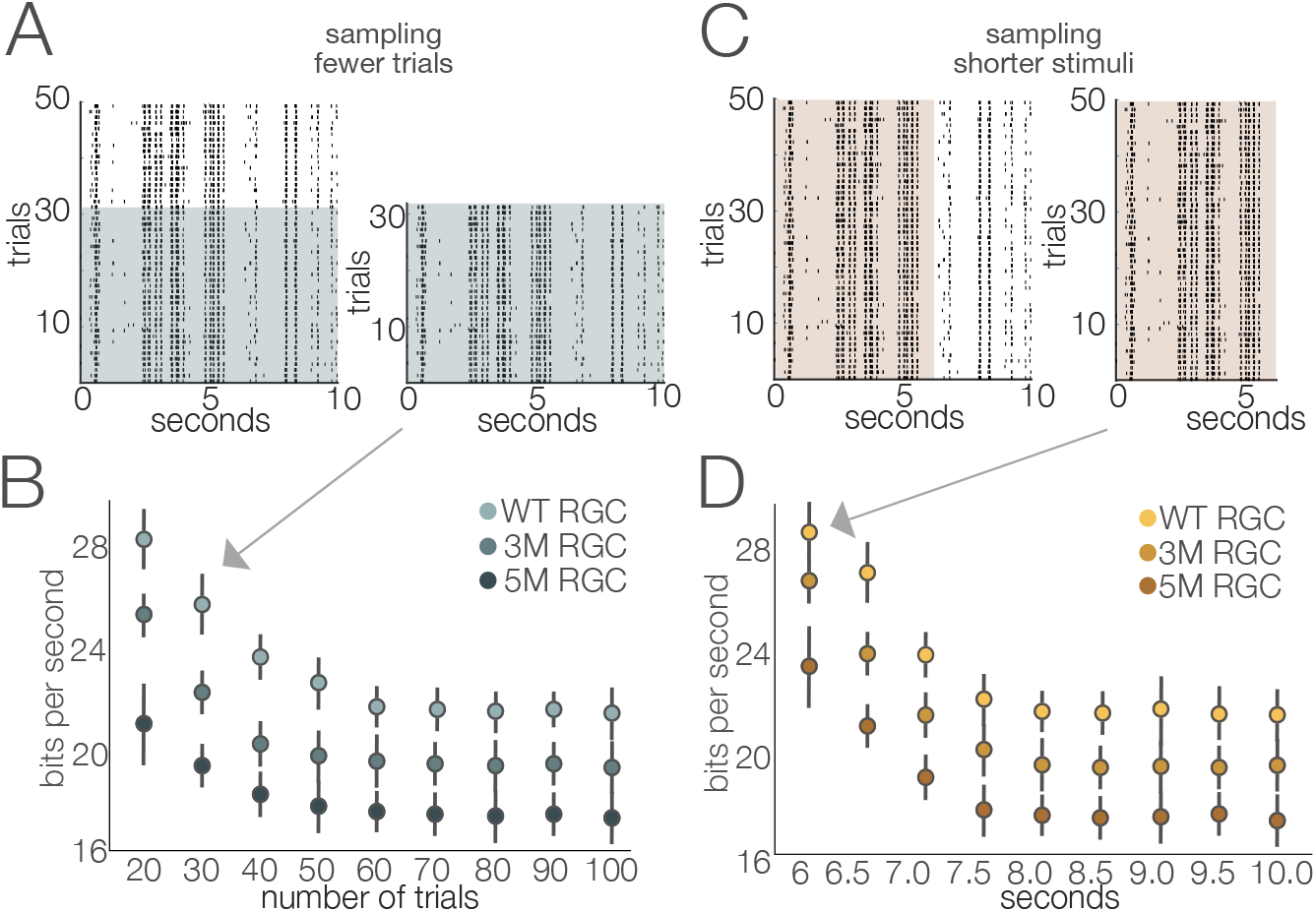
Stability of estimated information rates to trial number and trial duration. A. Example rasters illustrating the sub-sampling for fewer trials. B. Information rates (mean +/- 2SE) of 4 RGCs from separate cohorts as a function of number of repeated stimuli. Information rates stabilized at ∼60 trials (repeats). C. Example rasters illustrating the sub-sampling of trial duration. D. Information rates (mean +/- 2SE) of 3 RGCs from separate degeneration cohorts by sub-sampling briefer stimulus durations. The stimulus used in this analysis was movie 1. Similar results were obtained for the other stimuli (movie 2 and checkerboard noise).

### Visual compensation for cell loss begins in the retina

Human patients with RP45 (*Cngb1* mutations) maintain cone vision for many years, but the assumption has been that vision was preserved primarily through cortical compensation (Ferreira et al., 2017; Hickmott & Merzenich, 2002; Keck et al., 2013, 2008, 2011; Merabet & Pascual-Leone, 2010; Turrigiano, 2012). Indeed, studies in primary visual cortex using the *S334ter* rat model of RP support the notion that cortical plasticity bolsters visual processing during retinal degeneration (Chen et al., 2016). However, our findings indicate retinal signaling remains relatively robust under cone-mediated conditions despite large-scale rod loss, changes in cone morphology, and cone density. This suggests that at least some of the plasticity and homeostatic mechanisms required for vision during photoreceptor degeneration occur in the retina.

While we did not investigate mechanisms underlying this robust signaling, there is evidence that late-stage degenerated retinas maintain some photoreceptor to bipolar cell synapses (Beier et al., 2017; Beier, Palanker, & Sher, 2018; Care et al., 2019; Leinonen et al., 2020; Shen et al., 2020). Additionally, cone-mediated light responses are enhanced in a model with 50% rod loss compared to controls (Care et al., 2020). Mechanisms like these may help to explain why cone-mediated signaling was relatively stable until the latest stages of degeneration.

### Oscillatory spontaneous activity appears late in *Cngb1^neo/neo^* mice

A prominent feature of RP models is the emergence of abnormal spontaneous activity among RGCs (Trenholm & Awatramani, 2015). Previous studies on other models of RP, particularly *rd1* and *rd10* mouse models of Pde6b-RP, have shown abnormal spontaneous activity in two frequency bands: 1-2 Hz and 5-10 Hz (Biswas et al., 2014; Marc et al., 2007; Margolis et al., 2008; Menzler & Zeck, 2011; Poria & Dhingra, 2015; Stasheff et al., 2011; Tu, Chen, McQuiston, Chiao, & Chen, 2016; Ye & Goo, 2007). Abnormal horizontal cell activity is the source of the 1-2 Hz oscillations (Haq, Arango-Gonzalez, Zrenner, Euler, & Schubert, 2014), while AII amacrine cells are the likely source of the higher frequency oscillations (Borowska, Trenholm, & Awatramani, 2011; Choi et al., 2014; Ivanova, Yee, Baldoni, & Sagdullaev, 2016). Elevated and/or oscillatory spontaneous activity is a cause for concern in retinal degeneration because it is noise that competes with and deteriorates visual signals, particularly near threshold. It is unclear whether RGC oscillations also develop in humans with RP, although reports of phosphenes in RP patients could be explained by spontaneous activity (Gekeler, Messias, Ottinger, Ulrich Bartz-Schmidt, & Zrenner, 2006).

While previous studies have observed the emergence of oscillations in rodent models of RP (e.g., *rd10*) prior to the loss of photoreceptors and visual signaling, we only found evidence of spontaneous activity following total loss of photoreceptor signals in *Cngb1^neo/neo^* mice (9M, Figure 7). At 9M, we could not elicit visual responses from retinas when the video display was switched from off (0 Rh*/rod/s) to on with a ‘white’ screen (20,000 Rh*/rod/s). Furthermore, at 9M, there was no clear outer nuclear layer and cells that labeled for cone arrestin were very sparse and dysmorphic and did not co-label for M-opsin, indicating near total photoreceptor loss (data not shown). Even at 7M, when nearly all the rods and ∼30% of the cones have died, we observed no evidence for changes in spontaneous activity or oscillations. This is a promising observation from the perspective of gene therapy for rescuing photoreceptors for RP45: it suggests that elevated spontaneous activity and oscillation in RGC spiking will be avoided given that gene therapy is delivered prior to total cell loss.

### Implications for therapy

Our findings on the longevity of cone vision are particularly useful in determining the time window for therapeutic intervention in patients with RP45. Both pre-clinical and clinical studies have shown that late intervention does not ultimately halt photoreceptor cell death despite improvements to visually guided behaviors (Bainbridge et al., 2015; Cideciyan et al., 2013; Gardiner et al., 2020; Jacobson et al., 2015; S. Koch et al., 2012). It is currently unclear if the continued degeneration is slowed by therapy or not, or what the long-term implications are for vision restoration. Our study suggests that preserving cone-mediated vision in RP is entirely feasible considering that cone-mediated signals among RGCs are minimally perturbed by massive rod loss. However, it is not clear that rod-directed gene therapy will be successful at preserving cone (or rod) vision if it is delivered at a timepoint after which most of the rods have died. Thus, an important direction for future work is to determine at what timepoints rod-directed gene therapies need to be delivered to halt rod death such that cone vision can continue to function normally throughout the remaining lifespan of the treated individual.

## Materials and Methods

### Mice

Mice were used according to Duke University Institutional Animal Care and Use Committee guidelines and the Association for Research in Vision and Ophthalmology guidelines for the use of animals in vision research. All mice were housed with 12h light/dark cycles and fed rodent chow ad libitum. *Cngb1^neo/neo^*mice of both sexes were used (12 male and 12 female). The *Cngb1^neo/neo^* line has a neomycin resistance cassette inserted at intron 19 to disrupt Cngb1 mRNA splicing (Chen et al., 2010; Wang et al., 2019). Control animals (WT) consisted of male and female heterozygous littermates between 1 and 9M postnatal age to account for aging (4 male and 3 female). No age-related differences were found between WT mice (data not shown), so results were pooled. Genotyping was performed by Transnetyx using the primers/probe for the neomycin insert FWD GGGCGCCCGGTTCTT, REV CCTCGTCCTGCAGTTCATTCA, PROBE ACCTGTCCGGTGCCC, and primers/probe for WT Cngb1 FWD TCCTTAGGCTCTGCTGGAAGA, REV CAGAGGATGAACAAGAGACAGGAA, PROBE CTGAGCTGGGTAATGTC. At least 3 mice were used per timepoint, except to assay spontaneous activity at 9M, in which 2 mice were used (Figure 7).

### Immunohistochemistry and Confocal Microscopy

The sample of retina placed on the MEA and retina from the contralateral (unrecorded) eye were fixed for 30 minutes in 4% PFA (Thermo, 28908) at room temperature. Fixed eyes were hemisected with cornea and lens removed prior to immunolabeling.

For cryosections (Figure 1B-C), eye cups were placed in cold 30% sucrose for 3-12 hours, coated in Optimal Cutting Temperature Media (OCT; Tissue-Tek, 4583), placed in a microcentrifuge tube filled with more OCT, frozen using a bath of dry ice and 95% ethanol, and stored at −20°C for at least 24h. 12 µm sections were cut using a Leica cryostat (CM3050) and mounted onto frost free slides (VWR, 48311-703) stored at −20°C. To stain cryosections, slides were warmed to room temperature, rinsed 3x with 1x phosphate buffered saline (PBS; Santa Cruz, sc-296028), then incubated sequentially with 0.5% TritonX-100 (Sigma, X100) and 1% BSA (VWR, 0332) for 1h each. Primary antibodies were diluted with 0.3% TritonX-100 + 1% BSA, applied to slides at 4°C and incubated overnight. Slides were rinsed 3x with 1x PBS before applying secondary antibodies diluted with 1x PBS. After incubating at room temperature for 1h, slides were again rinsed 3x with 1x PBS, covered with mounting media containing DAPI (Invitrogen, P36935), cover slipped and sealed with clear nail polish.

For whole mounts (Figure 1D), retinal pieces were incubated with 4% normal donkey serum (NDS, Jackson Immuno, C840D36) in 1x PBS for >12h. Primary antibodies were diluted in 4% NDS. Tubes were shielded from light and placed on a rocker at 4°C for 7 days. Retinas were rinsed 3x with 1x PBS, then secondary antibody diluted in 1xPBS applied. Tubes were again placed a 4°C rocker for 1 day, rinsed 3x, mounted onto filter paper, mounting media applied, coverslipped, and sealed with nail polish.

All slides were kept at 4°C until imaged. Z-stack confocal images were taken using a Nikon AR1 microscope using 20x air and 60x oil objectives and motorized stage. Images were processed using FIJI software (Schindelin et al., 2012). Z-stacks were flattened according to their standard deviation. Brightness and contrast were adjusted as necessary. Cones were manually counted from 60x cross-sections labeled with antibodies to cone arrestin. Rod counts were obtained by measuring the area of the ONL and average DAPI nucleus count using the ‘Measure’ function in FIJI, with cone counts subtracted.

Antibodies:

Rabbit anti-mCar 1:500 (Millipore, AB15282)

Mouse anti-PCP2 1:500 (Santa Cruz, sc-137064)

Rabbit anti-M-opsin 1:500 (Sigma, AB5405)

Alexa-fluor secondaries 1:500 (Invitrogen, A31571 & A31572)

### MEA Recordings

Mice were dark adapted overnight by placing their home cage in a light shielded box fitted with an air pump for circulation. All dissection procedures the day of the experiment were carried out in complete darkness using infrared converters and cameras. Mice were decapitated, eyes enucleated and placed into oxygenated room-temperature Ames solution (Sigma, A1420) during retinal dissection and vitrectomy as described previously (Yao et al., 2018). A ∼1×2 mm piece from dorsal retina was placed RGC side down on a MEA with either 512 electrodes spaced 60-um apart, or 519 electrodes with 30-um spacing (Field et al., 2010; Frechette et al., 2005; Ravi, Ahn, Greschner, Chichilnisky, & Field, 2018). Oxygenated Ames perfused the retina throughout the experiment at a rate of 6-8 mL/min, heated to 32°C.

### Spike Sorting

Raw voltage traces from the MEA were spike sorted using custom software followed by manual curation as described previously (Field et al., 2007; Shlens et al., 2006). Cells with a spike rate >0.1 Hz and with <10% contamination estimated from refractory period violations were retained for further analysis.

### Visual Stimuli

The image from a gamma-calibrated OLED display (Emagin, SVGA + XL Rev3) was focused onto photoreceptors using an inverted microscope (Nikon, Ti-E) and 4x objective (Nikon, CFI Super Fluor ×4). Checkerboard stimuli were created and presented using custom Matlab code. Light from the OLED display was attenuated using neutral density filters. Retina was allowed to settle in darkness for approximately 30 min prior to initiating data collection. Then, spontaneous activity in darkness was recorded for 30 min. The display was then switched to a grey screen emitting ∼100 Rh*/rod/s and the retina was left to adapt for 5 min, after which mesopic spontaneous activity was recorded for 30 min. Repeated checkerboard noise was presented 200x, repeating every 10s. Additionally, non-repeating checkerboard noise was presented for 30min to estimate spatial and temporal RFs. For all mesopic checkerboard stimuli, the stimulus refreshed every 66 ms and the size of each square was 150×150 µm. After mesopic stimuli were presented, a static screen was presented and the NDF filter removed to allow the retina to adapt to a photopic light level (∼10,000 Rh*/rod/ s). Checkerboard noise repeats (200x, 10s) were presented, followed by 30 minutes of non-repeated checkerboard stimuli. For the photopic checkerboard movies, the stimulus refreshed every 33 ms and each square was 75×75 µm. Two 10s movie clips were presented 100x to estimate the mutual information between RGC signals and naturalistic movies. Movie 1 was a black and white video from a camera attached to a cat walking in the woods (Betsch, Einhäuser, Körding, & König, 2004). Movie 2 was a black and white video from a camera carried by a squirrel, depicting fast moving tree leaves (modified from Freiheit, 2016).

### Measuring receptive fields

RGC responses to checkerboard noise were used to estimate the spatial and temporal components of the spike triggered average (STA) (Chichilnisky, 2001). The STA estimates the spatial and temporal integration of visual signals within the RF of an RGC. For many RGCs, the STA is not space-time separable, meaning it cannot be expressed as the outer product of a function that depends only on space and a function that depends only on time. This precludes separately analyzing changes in spatial or temporal integration. To identify RGCs with space-time separable RFs, singular value decomposition (SVD) was performed on each STA. RGCs with STAs that were well-approximated by a rank-one factorization were kept for further analysis of their spatial and temporal RFs (Figure 2): these were cells for which the rank-one approximation captured >60% of the variance in the STA. For quantifying temporal integration, the time-to-zero crossing was used. This approximates the time-to-peak in the spiking response for a step increment (for ON RGCs) or step decrement (for OFF RGCs) of light (Chichilnisky & Kalmar, 2002). The analysis required that the temporal filter identified by SVD was biphasic because a well-defined zero-crossing was needed to estimate the time-to-zero: 79% of space-time separable STAs met this criterion.

### Mutual information

Mutual information (MI) was used to assess the fidelity of RGC signaling. MI measures how much observing the spike train reduces uncertainty about the stimulus. MI was estimated using the ‘direct method’ (Strong, Koberle, De Ruyter Van Steveninck, & Bialek, 1998). MI between a response and stimulus was computed as

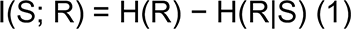

I(S;R) is the mutual information about the stimulus contained in the response. H(R) here is defined as the Shannon entropy of the response distribution (Shannon, 1948). This measure estimates the capacity of a neuron to convey information about the stimulus space. H(R|S) is the conditional entropy, a measurement of how noisy neural responses are across repeated trials of an individual stimulus.

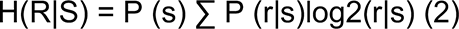

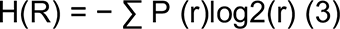

P(r) is the probability of a spike count occurring in a pattern across all trials of the stimulus space, while P(r|s) is the probability of observing a response pattern when a stimulus is presented. These probabilities are estimated by measuring the proportion of the observed response patterns at a given epoch of time from all the response patterns across all epochs of time.

Responses from 100 repeated trials of 10s white noise were used to estimate mutual information across all recorded RGCs. For each RGC, spike trains were binned to a time resolution that achieved the entropy estimates from the Ma upper bound (Ma, 1981; Strong et al., 1998). The response pattern length (a.k.a. ‘word’ length) was selected using the same procedure. The mutual information was calculated using the ‘direct method’ (Buračas, Zador, DeWeese, & Albright, 1998; Strong et al., 1998). Bins that achieved the Ma upper bound ranged from 4-6 milliseconds and response patterns ranged from 3-6 bins across all RGCs. The mutual information rate was computed as the quotient of the mutual information and the time length of the response pattern. Analysis of trial number and trial duration indicated that both were sufficiently large to produce stable estimates of the information rate (Figure 11).

#### Light Responsive RGCs

The proportion of RGCs that were light responsive was determined by computing a ratio between the variance and the mean in the peristimulus time histogram from the responses to the natural movie stimuli. The distribution of this ratio in an experiment was bimodal with unresponsive and responsive RGCs falling in each mode. A threshold was applied to each experiment to exclude the unresponsive RGCs.

### Spontaneous activity and power spectral analysis

The frequency spectra of spontaneous activity were calculated using the fast Fourier transform (FFT) applied to spike times binned at 1 ms from 30 min of spontaneous activity. Spectra were analyzed using a frequency range from 0.1 to 35 Hz. Peaks in the spectra were quantified for every RGC by computing: *abs(b-a)/abs(a-b)* where *b* represents the power at a baseline frequency level and *a* represents the maximum power. The baseline level was estimated by finding the average power in the 0.1 to 2 Hz range because this range was consistently flat across experiments and cells.

### Statistical tests

Significant changes across all assays were assessed using the two-way Kolmogorov–Smirnov test, a non-parametric test used to determine whether two sets of samples arise from the same distribution (Massey, 1952). P-values were corrected for multiple comparisons by Bonferroni correction. Error bars indicate the range of values within 2 standard errors (SE) of the mean, which was estimated by bootstrapping the mean 2000 times.

To measure whether differences across time points could be produced by other factors (e.g., experiment-to-experiment variability), a parametric linear mixed effects model was used (Bates, Mächler, Bolker, & Walker, 2015). The mixed effects model accounts for retina-to-retina variability by adding each experiment as a random effect. This procedure permitted making broad level inferences about the RGC populations without dependence on experimental variability. In addition, the sex of the animal was considered by including it as an interaction term with the degeneration conditions. This step enabled determining whether degeneration conditions were associated with information rates and RF sizes in a sex-independent fashion. The model indicated that conclusions about the impact of degeneration on RGC signaling were insensitive to both sex and experiment-to-experiment variability.

## Acknowledgements

We thank our funding sources: National Institute of Health (NIH) R01 EY024280-01, NIH NEI core grant EY5722, Holland Trice Foundation, Whitehead Foundation, and Research to Prevent Blindness unrestricted grant to Duke University. We also are grateful to Dr. Jon Cafaro and Dr. Suva Roy for technical assistance and Erika Ellis for discussions.

